# An ancient, conserved gene regulatory network led to the rise of oral venom systems

**DOI:** 10.1101/2020.08.06.240747

**Authors:** Agneesh Barua, Alexander S. Mikheyev

## Abstract

Oral venom systems evolved multiple times in numerous vertebrates enabling exploitation of unique predatory niches. Yet how and when they evolved remains poorly understood. Up to now, most research on venom evolution has focussed strictly on the toxins. However, using toxins present in modern day animals to trace the origin of the venom system is difficult, since they tend to evolve rapidly, show complex patterns of expression, and were incorporated into the venom arsenal relatively recently. Here we focus on gene regulatory networks associated with the production of toxins in snakes, rather than the toxins themselves. We found that overall venom gland gene expression was surprisingly well conserved when compared to salivary glands of other amniotes. We characterised the ‘meta-venom’, a network of approximately 3000 non-secreted housekeeping genes that are strongly co-expressed with the toxins, and are primarily involved in protein folding and modification. Conserved across amniotes, this network was co-opted for venom evolution by exaptation of existing members and the recruitment of new toxin genes. For instance, starting from this common molecular foundation, *Heloderma* lizards, shrews, and solenodon, evolved venoms in parallel by overexpression of kallikreins, which were common in ancestral saliva and induce vasodilation when injected, causing circulatory shock. Derived venoms, such as those of snakes, incorporated novel toxins, though still rely on hypotension for prey immobilization. These similarities suggest repeated co-option of shared molecular machinery for the evolution of oral venom in mammals and reptiles, blurring the line between truly venomous animals and their ancestors.

## Introduction

Venoms are proteinaceous cocktails that can be traced and quantified to distinct genomic loci, providing a level of genetic tractability that is rare in other traits(*1*–*4*) This advantage of venom systems provided insights into processes of molecular evolution that are otherwise difficult to obtain. For example, studies in cnidarians showed that gene duplication is an effective way to increase protein dosage in tissues where different ecological roles can give rise to different patterns of gene expression(*2*, *5*). Studies of venom in snakes have allowed comparisons of the relative importance of sequence evolution vs gene expression evolution, as well as how a lack of genetic constraint enables diversity in complex traits(*6*, *7*).

Despite the wealth of knowledge venoms have provided about general evolutionary processes, the common molecular basis for the evolution of venom systems themselves is unknown. Even in snakes, which have perhaps the best studied venom systems, very little is known about the molecular architecture of these systems at their origin(*8*, *9*). Using toxin families present in modern snakes to understand evolution at its origin is difficult because toxins evolve rapidly, both in terms of sequence and gene expression(*10*, *11*). Toxins experience varying degrees of selection and drift, complicating interpretations of evolutionary models(*12*), and estimation of gene family evolution is often inconsistent, varying with which part of the gene (exon or intron) is used to construct the phylogeny(*13*). Most importantly, present-day toxins became a part of the venom over time, this diminishes their utility in trying to understand events that lead to the rise of venom systems in the ancestors of snakes(*14*, *15*).

A gene co-expression network aims to identify genes that interact with one another based on common expression profiles(*16*). Groups of co-expressed genes that have similar expression patterns across samples are identified using hierarchical clustering and are placed in gene ‘modules’(*17*). Constructing a network and comparing expression profiles of modules across taxa can identify key drivers of phenotypic change, as well as aid in identifying initial genetic targets of natural selection(*18*, *19*). Comparative analysis using gene co-expression networks allows us to distinguish between ancient genetic modules representing core cellular processes, evolving modules that give rise to lineage-specific differences, and highly flexible modules that have evolved differently in different taxa(*20*). Gene co-expression networks are also widely used to construct gene regulatory networks (GRN) owing to their reliability in capturing biologically relevant interactions between genes, as well as their high power in reproducing known protein-protein interactions(*21*, *22*).

Here we focus on gene co-expression networks involved in the production of snake venom, rather than the venom toxins themselves. Using a co-expression network, we characterized the genes associated with venom production, which we term the ‘meta-venom’, and determine its biological role. We traced the origin of this network to the common ancestor of amniotes, which suggests that the venom system originated from a conserved gene regulatory network. The conserved nature of the meta-venom across amniotes suggests that oral venom systems likely started with a common gene regulatory foundation, and underwent lineage-specific changes to give rise to diverse venom systems in snakes, lizards, and even mammals.

## Results

### The meta-venom is associated with toxin expression in the venom gland of snakes

Previously published RNA libraries from Taiwan habu (*Protobothrops mucrosquamatus*) were used to construct the network(*12*). Weighted Gene Co-expression Network Analysis (WGCNA) was used to construct the co-expression network(*23*). WGCNA estimates correlations between genes across samples (libraries) and clusters genes with similar profiles into modules(*23*).

Using data from venom gland samples WGCNA clustered 18,313 genes into 29 modules ranging in size from 38 to 3380 genes. All secreted venom toxins were found in the largest module (module 1), which we term the ‘meta-venom’ (Fig 1a). Therefore, the meta-venom represents an assemblage of housekeeping genes that are strongly associated with toxin genes. These form an ensemble that is the gene regulatory network (GRN) involved in expression of toxin genes. We performed module preservation analysis to determine whether within-module characteristics like gene density and connectivity between genes are conserved between venom gland and other tissues like heart, kidney, liver. In other words, module preservation statistics were used to determine whether the characteristics of genes and their modules identified in one (reference) tissue were present in another (test) tissue. A module preservation Z_summary_ > 2 implies that module characteristics within a module are preserved in other tissues, while a score < 2 denotes no preservation(*24*). Z_summary_ statistic (Supplementary table 1) revealed that the meta-venom module is not preserved in the heart or liver, but has borderline preservation in the kidney (Z_summary_ = 2.000522). This implies that much of the expression pattern of the meta-venom is unique to the venom gland and bears only a slight similarity in kidneys.

**Fig 1:**
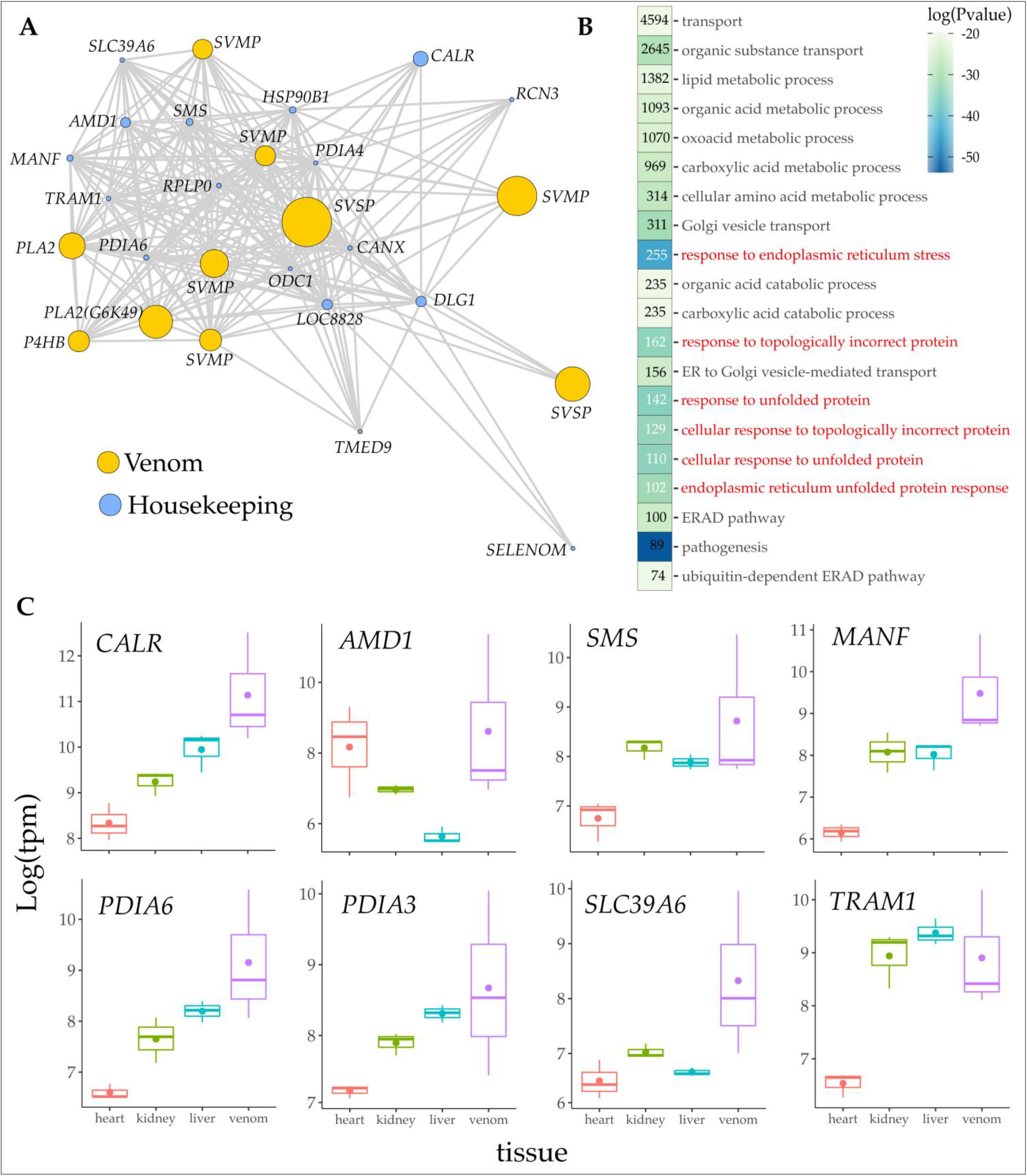
The meta-venom module represents a group of co-expressed genes that are associated with production of toxin in the venom gland of the Taiwan habu. More than one-third genes in the meta-venom are upregulated in the venom gland, and are involved in protein folding and protein modification. **a)** The meta-venom module comprised a total of 3380 genes. Out of them 10 of the most expressed toxin genes and 20 of the most expressed non-venom were plotted to visualize connections and overall module topography. An interactive version of the network graph is available at (Supplementary figure 1). Most toxin genes and non-toxin genes are well interconnected. ( LOC8828 represents a gene without a reliable annotation, but we believe it is a truncated SVMP and it is flanked very closely by a secreted SVMP, however, since it is not secreted in the venom we classify it as a housekeeping gene) b) The 20 most significant GO terms enriched in the meta-venom module comprised processes related to molecular transport and metabolism. We focussed on the most significantly enriched GO terms (in red) as they represent more specific biological processes and are less ambiguous as compared to more broadly defined terms like ‘transport’ and ‘organic acid metabolic process’. These specific terms refer to processes involved in protein folding and modification, in particular; the unfolding protein response (UPR), and endoplasmic reticulum associated protein degradation (ERAD). The GO term ‘pathogenesis’ has the highest significance and is attributed to the toxin-genes present in the meta-venom. GO terms are arranged by descending order of size (given within panels). **c)** Most of the genes with high module membership were on average upregulated in the venom gland, with some upregulated in non-venom tissue. *CALR:* calreticulin*, AMD1:* adenosylmethionine decarboxylase 1*, SMS:* spermine synthase*, MANF:* mesencephalic astrocyte derived neurotrophic factor*, PDIA6:* protein disulfide isomerase family A member 6*, PDIA3:* protein disulfide isomerase family A member 3*, SLC39A6:* solute carrier family 39 member 6*, TRAM1:* translocation associated membrane protein 1. Therefore, the UPR and ERAD pathway seem particularly important for venom expression and likely helps maintain tissue homeostasis under the load of high protein secretion.

After defining the meta-venom, which comprises genes that are tightly associated with toxin expression, we identified the biological processes involved using Gene Ontology (GO) enrichment. The meta-venom is primarily involved in protein modification, and protein transport (Supplementary table 2). GO terms associated with the unfolding protein response (UPR): *GO:0006986, GO:0034620, GO:0035966*, and endoplasmic reticulum associated protein degradation (ERAD): *GO:0034976, GO:0030968, GO:0036503* were the most significantly enriched biological processes in the meta venom (Fig 1b).

Since the meta-venom has over 3000 genes, visualising the entire network topology would be impractical. Therefore, we selected the top 20 highly expressed non-venom genes, and the top 10 highly expressed toxin genes for visualization and to identify the levels of connection between them (Fig 1a). An interactive visualization can be found in the online supplementary material (Supplementary figure 1). The network diagram revealed that almost all of the highly-expressed venom toxins interact strongly with each other, as well as directly with the non-venom genes. Zinc metalloproteinase (SVMP: 107298299) and snake venom serine protease serpentokallikrein-2 (SVSP: 107287553) were the exceptions, which interact with only a few toxin genes and non-venom genes (namely DLG1, CANX, HSP90, RPLP0, PDIA4, and LOC8828).

Several network characteristics can be used to identify genes integral to a network. One of these characteristics is module membership, which represents connectivity of a gene with other genes within a module and is used to define centralised hub genes(*23*). Module membership (MM) has values between 0 and 1, where values closer to 1 represent high connectivity within a module, and values closer to 0 represent low connectivity. We estimated module membership of genes in the meta-venom and identified sets of differentially expressed genes (DEGs) (Supplementary table 3). An ANOVA-like test for gene expression in venom gland, heart, liver, and kidney of habu revealed that out of 3380 genes that make up the meta-venom, 1295 were significantly differentially expressed (*p* < 0.05) (Supplementary table 3). To identify genes most specific to the venom gland we filtered the DEGs associated with the UPR and ERAD that had high module membership (MM > 0.9) and high average expression across all venom gland libraries. We obtained a list of 149 genes (Supplementary table 3). On an average, most of these genes were upregulated in the venom gland, with a few upregulated in the non-venom tissues (Fig 1c, only 8 shown, full data set in online Supplementary table 3), implying that these genes are of greater functional relevance in the venom gland.

#### External validation of module preservation

To confirm that modules identified in this study, particularly the meta-venom module, represent technically reproducible and evolutionarily meaningful features, we assessed the extent of module preservation between our work and a WGCNA investigation of the human salivary gland (*25*). Other than the WGCNA algorithm, this study employed different methodologies, such as microarray gene expression measurements, and the inclusion of samples from patients with salivary gland pathogenesis. Nonetheless, there were significant overlaps in modules detected in both studies, supporting the method’s robustness (Supplementary figure 2).

### The meta-venom is conserved across amniotes

Conserved gene expression profiles between taxa are indicative of a shared ancestry that can be used to provide insights into key drivers of phenotypic change as well as revealing molecular organization of a trait at its origin(*17*, *20*). The meta-venom is significantly enriched for genes belonging to the UPR and ERAD pathways. These families of housekeeping genes are widely conserved across the animal kingdom(*26*). This high level of conservation encouraged the search for orthologs in other taxa. Once the list of orthologs was obtained we carried out comparative transcriptomic analysis to determine if the expression of meta-venom were conserved across taxa. We identified 546 one-to-one orthologs of the meta-venom, that were expressed in 4 tissue groups of 9 species: human, chimpanzee, mouse, dog, anole, habu, cobra, chicken, and frog. To do this we first obtained one-to-one orthologs from NCBI’s eukaryotic genome annotation pipeline and combined them with phylogenetically inferred orthologs from *OrthoFinder(27, 28)*. In addition to the substantial overlap between estimated orthologs, both approaches estimated orthologs with conserved synteny (Supplementary figure 3). Public RNA datasets from 4 tissues (heart, kidney, liver, and salivary glands) were used for comparative transcriptomic analysis (see methods). We obtained expression data for cobra tissues, including that of venom gland from(*29*).

**Fig 3:**
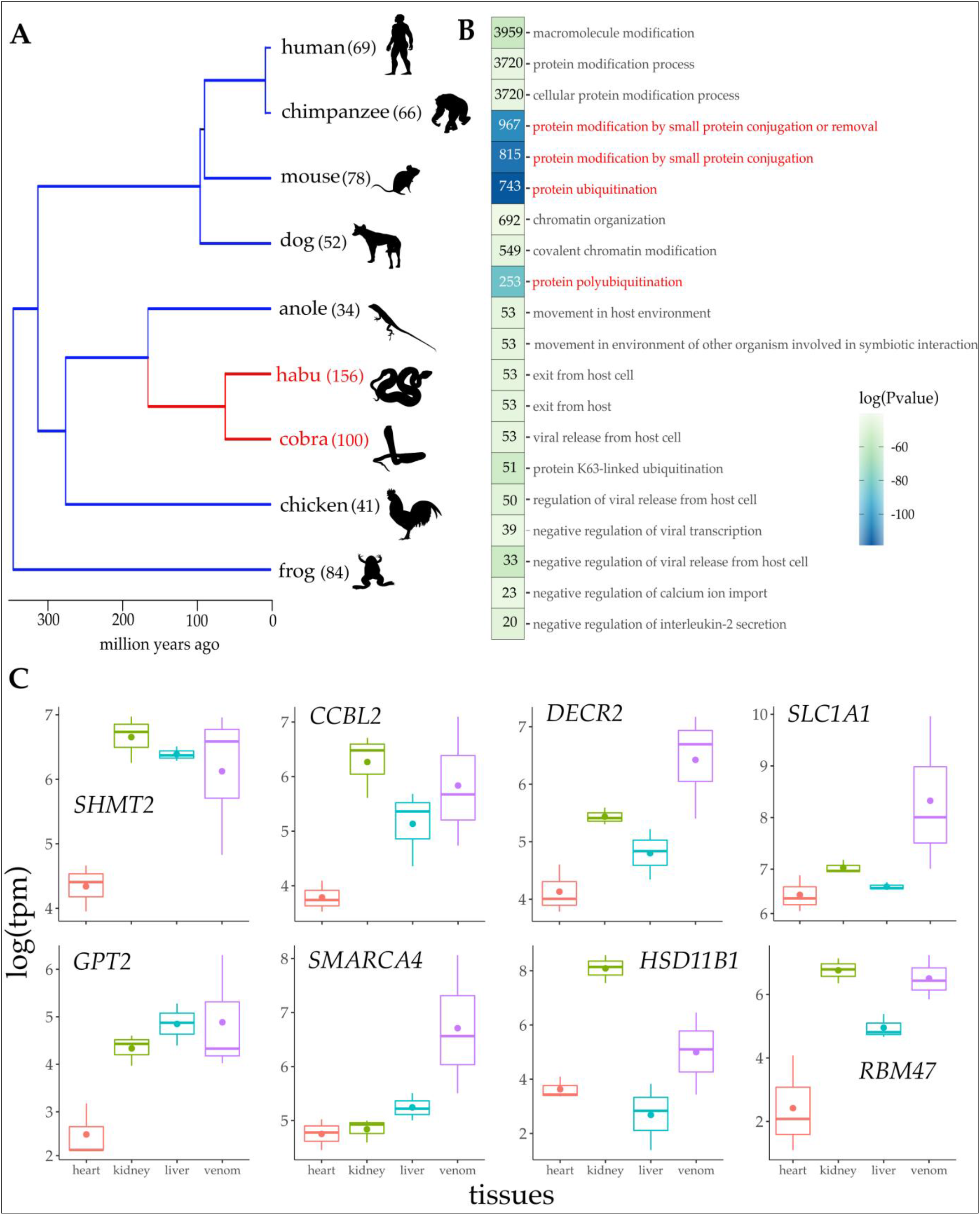
Gene families in the meta-venom have not only evolved more rapidly in the lineage leading to snakes, but have also undergone more expansions in snakes than in other taxa. **a)** Gene family evolution modeled as a ‘birth and death’ process revealed higher rates of evolution in the branch leading up to venomous snakes (red; λ = 6.4 × 10^−3^) as compared to other taxa (blue; λ = 1.7 × 10^−3^). A model with dual rates (λ1, λ2) at different branches was a better fit than a uniform rate (single λ across the whole tree) model as estimated by a likelihood ratio test (Supplementary figure M5). **b)** Orthogroups undergoing significant expansion were highly enriched for GO terms protein ubiquitination (*GO:0016567*), protein modification by small protein conjugation (*GO:0032446*), protein modification by small protein conjugation or removal (*GO:0070647*), and protein polyubiquitination (*GO:0000209*), among others. **c)** On average, most of the genes that were associated with the above GO terms, were upregulated in the venom gland, although a substantial portion was upregulated in other tissues as well (only 8 shown, full list in Supplementary table 2). *SHMT2:* serine hydroxymethyltransferase 2 (mitochondrial)*, CCBL2:* cysteine conjugate-beta lyase 2*, DECR2:* 24-dienoyl-CoA reductase 2 peroxisomal*, SLC1A1:* solute carrier family 1 member 1*, GPT2:* glutamic pyruvate transaminase (alanine aminotransferase) 2*, SMARCA4:* SWI/SNF related matrix associated actin dependent regulator of chromatin subfamily a member 4*, HSD11B1:* hydroxysteroid (11-beta) dehydrogenase 1*, RBM47:* RNA binding motif protein 47.

To get an overview of meta-venom gene expression patterns between species, we performed a principal component analysis (PCA) using a comparative data set of the one-to-one meta-venom orthologs. PCA clustered gene expression by tissue and despite the over 300 million years’ divergence between the taxa, differences among tissues explain more than 30% of variation present in the data (Fig 2a). Performing a PCA using all 2682 expressed orthologs between 9 taxa, including those outside the meta-venom, homologous tissues clustered more tightly (Supplementary figure 4). As a sanity check we chose orthologs at random to check whether the transcriptomes would still be clustered by tissue; however, a random set of genes produced no clustering (Supplementary figure 5). This indicated that tissues cluster together based on some underlying structure in the expression patterns of specific sets of genes analysed, and that this clustering cannot be reproduced by using any arbitrary set of genes(*30*). Our comparative transcriptomic analysis showed that expression patterns between homologous tissues were well conserved, especially between venom glands in snakes and salivary glands in mammals. This suggests that the gene regulatory architecture of the meta-venom likely evolved in the common ancestor of amniotes and has for the most part remained conserved in extant taxa, while giving rise to the venom gland in snakes.

**Fig 2:**
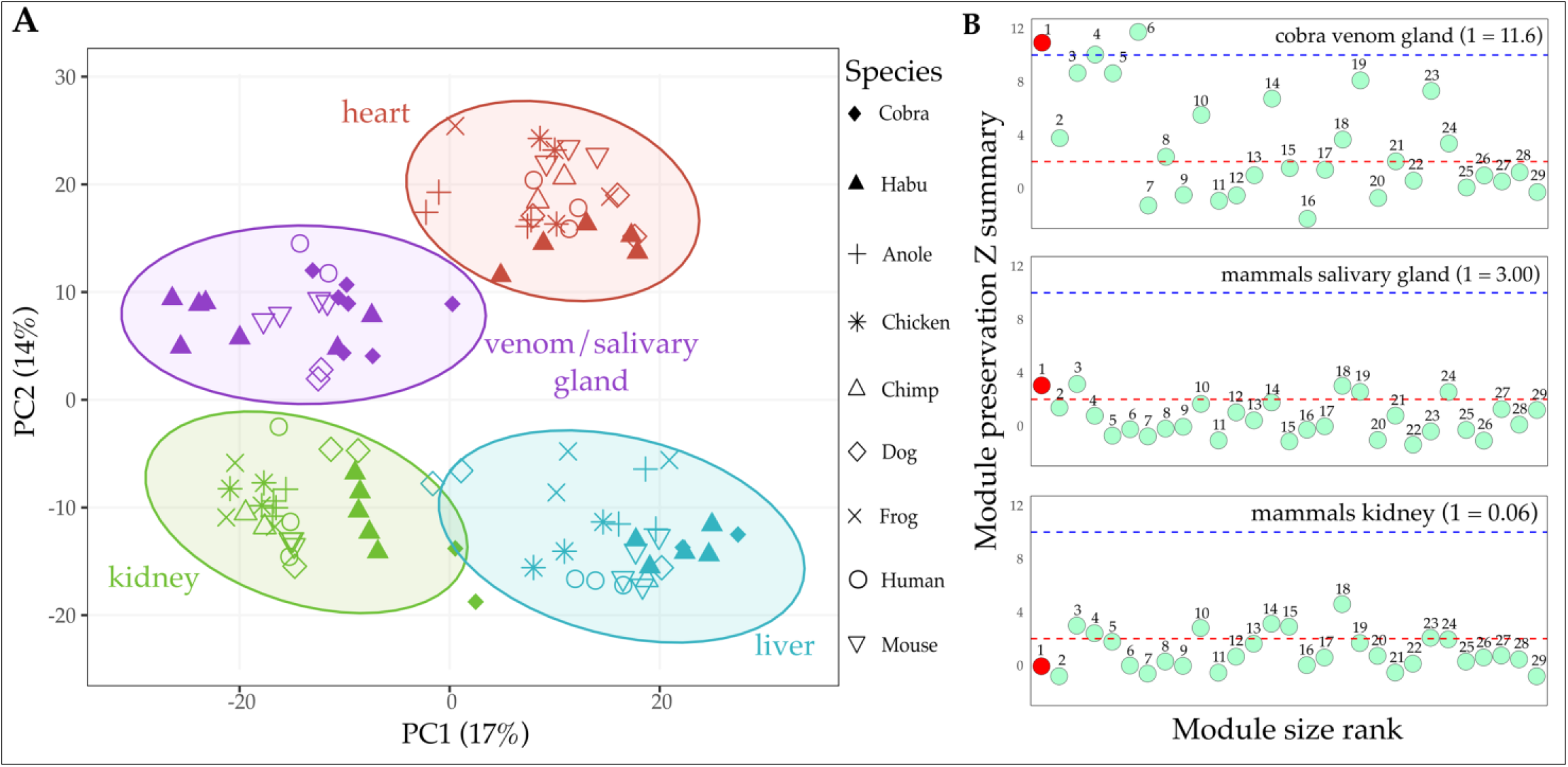
Expression pattern of orthologs between venom gland in snakes and salivary gland in mammals was surprisingly well conserved. This conservation was also reflected in the preservation of the meta-venom module in the salivary gland of mammals. **a)** When selecting the 546 one-to-one meta-venom orthologs expressed in all 9 species, transcriptomes clustered based on tissue. Ellipses represent 95% confidence intervals. GO term enrichment of these 546 genes revealed that genes from the UPR and ERAD pathway are still significantly enriched, suggesting that even in a reduced dataset, the functional core of the meta-venom is still conserved (Supplementary table 2). Despite the large evolutionary distance between species (MRCA ~ 300 million years ago), partitioning by tissue explains >30% of the variation in the data. **b)** The meta-venom was highly preserved in the venom gland of cobra (Z_summary_ > 10) while it was weakly preserved in salivary gland of mammals (Z_summary_ > 2). The meta-venom however, was not preserved in the kidneys of mammals (Z_summary_ < 2). These lines of evidence indicate that common regulatory architecture inherited from a common amniote ancestor gave rise to the snake venom gland. Despite the subsequent evolutionary elaboration of the venom gland, it has maintained this conserved regulatory core.

### Network characteristics of the meta-venom are conserved between the salivary glands of mammals and venom glands of snakes

The clustering of transcriptomes of venom gland in snakes and salivary gland in mammals was interesting because it suggests that both these tissues have a degree of molecular conservatism that likely originated in their common ancestor. Therefore, to test whether the modular characteristics of the meta-venom are preserved in the salivary tissue of mammals we carried out module preservation analysis.

We estimated module preservation of the meta-venom in the venom gland of cobra and the salivary tissue of several mammals where sufficient transcriptomic data were available (mouse, human, and dog). The meta-venom was preserved in both venom glands of cobra as well as salivary tissue of mammals (Fig 2b). In cobra, the meta-venom had a Z_summary_ > 10 implying very high preservation, while in salivary tissue of mammals the Z_summary_ was 3, implying weak to moderate preservation.

Despite the differences between these tissues however, the fact that the meta-venom is weak to moderately preserved (Z_summary_ > 2) in salivary tissues of mammals provides evidence of a degree of molecular conservatism that has likely persisted since the origin of oral secretory tissues in amniotes.

### Gene families in the meta-venom evolve rapidly and have undergone greater expansion in venomous snakes

Increasing the number of gene copies, especially in venom systems are crucial to bringing about evolutionary novelty(*2*, *31*, *32*). The meta-venom in habu comprises genes that have many copies, which could have played a role in evolution of the venom system in snakes (Supplementary table 4). To determine whether gene families in the meta-venom evolved rapidly in venomous snakes, either by expansions or contractions, we examined gene family evolution using CAFE(*33*).

We used different rate parameters (λ) along the lineage leading up to venomous snakes to test the hypothesis that meta-venom gene families evolved faster in snakes as compared to other species. The rate parameter λ describes the probability that any gene will be either gained or lost, where a higher λ denotes rapid gene family evolution(*34*). Gene families in the branches leading up to snakes have a higher degree of family expansion, as well as higher evolution rates (λ = 6.450 × 10^−3^) as compared to the rest of the tree (λ = 1.769 × 10^−3^) (Fig 3a). Among the orthogroups identified by CAFE, 23 groups were statistically rapid (see Methods). Ancestral estimations of gene family sizes showed that in the venomous snake lineage, most families (16 out of 23) underwent significant expansions, while a few families contracted (2 out of 23) or remained the same (5 out of 23) (Supplementary table 5). GO term enrichment of the 23 statistically rapid orthogroups revealed genes involved in protein modifications, protein ubiquitination, viral release from cells (genes from snakes, not of viral origin), and chromatin organization, among others (Fig 3). We focussed on genes having the most significant GO terms (Fig 3b), namely, protein ubiquitination (*GO:0016567*), protein modification by small protein conjugation (*GO:0032446*), protein modification by small protein conjugation or removal (*GO:0070647*), and protein polyubiquitination (*GO:0000209*). Of the expanded genes in the meta-venom that were enriched for these terms, almost half were significantly differentially expressed between venom gland, heart, liver, and kidney (Supplementary table 3). While on average, most of these genes were upregulated in the venom gland, many were upregulated in other tissues (Fig 3c, only 8 shown, full list in Supplementary table 3). Our results show that although genes involved in protein ubiquitination underwent significant expansion in venomous snakes, their overall activity is not strictly restricted to the venom gland but functions in other tissues as well.

## Discussion

No biological system acts in isolation, even highly specific processes. Co-expression of genes regulates both cellular processes and maintains cellular homoeostasis(*20*, *35*, *36*). Toxin genes in the snake venom system are co-expressed with a large number of non-toxin genes. Together they form a gene regulatory network (GRN) that we term the ‘meta-venom’. The meta-venom comprises genes that are involved in various processes, the most significant being the unfolded protein response (UPR) and endoplasmic reticulum associated protein degradation (ERAD) pathways. While toxin genes are evolutionarily labile(*37*), the conserved genes they interact with reveal the origins and repeated evolution of venom systems in vertebrates.

Genes with evolutionarily conserved expression represent functionally important groups in which co-regulation is advantageous(*20*). Therefore, the conserved expression of meta-venom orthologs between venom glands in snakes and salivary glands in mammals was particularly important (Fig 2a). While many snakes employ an oral venom system for securing prey, there are also mammals, such as shrews, and Solenodons, that have evolved oral venom systems (based on salivary glands) for prey capture or defence(*38*). Therefore, the overall conservation of meta-venom expression, as well as preservation of the meta-venom module (Fig 2b), suggests that salivary glands in mammals and venom glands in snakes share a functional core that was likely present in their common ancestor. Using this common molecular foundation as a starting point, snakes diversified their venom systems by recruiting a diverse array of toxins while mammals developed less complex venom systems with high similarity to saliva(*39*). Developing similar traits using common molecular building blocks is the hallmark of parallelism(*40*).

Despite the shared molecular foundation, however, the alternate path taken by snakes and the majority of mammals in developing an oral secretory system has led to the accumulation of large scale phenotypic and functional differences between the two lineages. For instance, salivary tissue of most mammals produce large volumes of very dilute mixtures, while snake venom glands produce highly concentrated mixtures of diverse toxins(*41*). At the genetic level these differences are apparent when comparing evolutionary rates of gene families that comprise the meta-venom. In venomous snakes, gene families have undergone greater expansions, and have evolved at a significantly higher rate than in other lineages like mammals (Fig 3a). The most enriched process among the groups of significantly expanded gene families is protein modification via ubiquitination (Fig 3b). Along with tagging proteins for degradation, the ubiquitin system influences various aspects of protein functioning in the cell(*42*). The significant expansion of these genes in venomous snakes suggests a possible link between establishment of a complex venom system and the need for a molecular machinery which shapes a multitude of cellular processes.

### The UPR and ERAD system likely promoted the evolution of an oral venom system

While it is difficult to attribute individual genes to a specific process without functional assays, knowing how the components of the meta-venom function in other species, we can hypothesise their roles in the venom gland of snakes and their ancestors. This helps paint a picture as to how incorporating these genes would enable the establishment of an oral venom system.

The UPR and ERAD act as ‘quality control’ machinery ensuring that proteins undergo proper folding and maturation(*43*). Several hub genes in the meta-venom that are upregulated in the venom gland can contribute to this quality control process (Fig 1c). For example, Calreticulin (CALR) is a lectin-like chaperone that increases both the rate and yield of correctly folded proteins as well as preventing aggregation of partially folded proteins(*44*). Mesencephalic astrocyte derived neurotrophic factor (MANF) is induced during the UPR as a response to overexpression of misfolding-prone proteins to alleviate ER stress, and has an evolutionarily conserved cytoprotective function(*45*, *46*). Disulfide bonds maintain structural stability and functional integrity of many secreted proteins including venom toxins(*47*). Our results confirmed this as protein disulfide isomerase families (*PDIA6, PDIA3*) were upregulated in the venom gland and also occupied hub positions in the meta-venom (Fig 1a, c). PDI families catalyse disulphide bond formation, and are also vital in rearranging incorrect bonds to restore correct protein conformation(*48*). It is perhaps this restorative ability of PDI that makes it integral to the meta-venom. Individual components of the UPR and ERAD also do not work in isolation. Feedback loops allow several components to communicate and coordinate their individual processes to relieve ER stress. For example, CALR and PDIA work in close association to equilibrate the removal of misfolded proteins and restore correct protein conformation(*49*). This is perhaps reflected in the meta-venom where they not only share connections, but also occupy hub positions.

Although UPR and ERAD are considered to be stress responses, they function in a stress-independent manner as well. The UPR system is activated by developmental, cell surface signalling, circadian, and various other physiological cues, implying that the system (or at least elements of it) are frequently and perhaps even continuously fine tuning cellular functions(*50*). In fact, consistent detection of key regulators of the UPR (ATF4, ATF6, and PERK) in non-stressed mouse tissues suggest their role in basal regulation of gene expression *in vivo(51–53)*. Having UPR regulators contribute to the regulation of various cellular processes provides greater flexibility: a wide range of signals can be transmitted to multiple overlapping or branching pathways to fine tune their activity, a form of regulation that would be evolutionarily advantageous in organisms with diverse tissue types(*50*). This fine tuning is further enhanced by ubiquitin ligases that spatially and temporally modify the magnitude and duration of the UPR, impacting overall physiology(*54*). Therefore, the expansion of meta-venom genes associated with protein ubiquitination (Fig 3c) likely enabled a high degree of fine tuning of cellular secretory processes in lineages leading up to venomous snakes.

The UPR anticipates, detects, and correctly folds misfolded proteins. The ERAD ensures that misfolded proteins are degraded so as to prevent cellular toxicity, and ubiquitin ligases add an overall level of regulation to fine tune these processes. Having such a robust regulatory network in place should improve the tenacity of the system, enabling it to tolerate an increase in tissue complexity through changes in composition and concentration of secreted proteins. Therefore, having these molecular systems already in place likely primed the ancestors of venomous animals to undergo a series of steps to attain a weaponised oral venom system.

### Evolution of oral venoms from an ancestral salivary GRN

Given the existence of a conserved salivary GRN, venom can evolve in two ways: exaptation of existing components or through the addition of novel genes. Both mechanisms played a role in the evolution of snake venom. Furthermore, the architecture of the ancestral salivary GRN and comparisons to other venoms, such as those of solenodon and shrews, suggests a general model by which venoms have evolved across a range of taxa:

### Stage 1: Exaptation of salivary enzymes, particularly kallikreins

Kallikreins are a group of trypsin or chymotrypsin-like serine proteases whose origins can be traced back to the common ancestor of tetrapods almost 330 million years ago(*55*). They are expressed in multiple tissues, and are especially abundant in saliva, suggesting that were likely an important component of the ancestral saliva(*55*, *56*). Kallikrein proteolytic activity releases bradykinin and promotes inflammation, and may be involved in wound healing(*57*). Interestingly, when injected, salivary kallikreins from non-venomous animals, such as mice and rats induce a hypotensive crisis leading to death(*58*, *59*). In fact, Hiramatsu *et al.(58)* effectively blurred the lines between venomous and non-venomous mammals by proposing that male mice secrete “toxic proteins (kallikrein-like enzymes) into saliva, as an effective weapon.” Lethality of saliva differs between mouse strains, suggesting that heritable variability in this trait exists within species(*60*), a necessary prerequisite for adaptation. Selection on increased lethality, particularly through higher levels of protein secretion, could have led to the evolution of other oral venoms in mammals (*e.g.*, solenodon and *Blarina* shrews), as well as *Heloderma* lizards, which all employ kallikrein overexpression(*39*, *61*, *62*). Similarly, Fry(*14*) noted that snake venom kallikreins most likely arose by direct modification of salivary counterparts, based on their phylogenetic proximity to salivary proteins in lizards. The high similarity between venom and salivary secretions in solenodon seems to confirm the saliva to venom path of toxin recruitment(*39*). Intriguingly, a recent report on caecilian venom detected fibrinogenolytic activity(*63*), which is also consistent with the presence of kallikreins(*64*). It appears that the ancestral salivary GRN’s composition predisposes the evolution of envenomation strategies based on hypotensive shock via kallikrein injection. Intriguingly, despite the complexity of the modern snake venom arsenal, hypotensive shock remains one of two main strategies for prey immobilization (*65*).

While kallikreins represent the most striking and taxonomically diverse example of exaptation, other ancestral salivary components have been recruited by a range of taxa. For instance, cysteine-rich secretory proteins (CRISPs), which are expressed in many tissues including salivary glands, are commonly found in the venom of snakes and of lizards (*Heloderma)(14, 66)*. CRISPs play a wide variety of roles in non-venomous tissues, and their function appears likewise diverse in venoms(*67*). Vampire bats use a calcitonin gene-related peptide (CGRP) as a vasodilator(*68*), and this peptide is likewise expressed in human and rat saliva, as well as other tissues(*69*, *70*). As CGRPs are potent vasodilators, their co-option into bat venom should enhance its function. These examples illustrate that the ancestral expression of a gene need not be limited to saliva since many of them are also expressed in other tissues as well, as are many, if not all, elements of the meta-venom (Figure 1c, 3c). Rather, these genes are united by pharmacology that could be easily re-purposed and overexpressed in the novel venomous context. It further suggests that the salivary GRN is flexible, in that it can evolve to secrete high levels of a wide range of proteins.

### Stage 2: gene recruitment

Snake venoms arose from the same ancestral GRN and likely followed the same first evolutionary step relying on initial exaptation of existing components. Yet, today they contain numerous novel toxins and bear little resemblance to the composition of ancestral saliva. Incorporation of novel toxins has occurred relatively infrequently, and the process remains poorly understood at the transcriptional level. For example, recent insights into the evolution of snake venom metalloproteinases found that they are related to the mammalian *adam28* gene(*32*, *71*). This gene is expressed in many tissues, but only weakly in the salivary glands of some species(*72*), and, furthermore, it is a transmembrane rather than a secreted protein. While the series of sequential deletions necessary for the protein sequence to acquire toxicity have been revealed(*32*), the corresponding changes in gene expression accompanying them remain a mystery. Similarly, while the origin of phospholipases A_2_ has been traced to a common amniote ancestor, the steps required for its neofunctionalization remain obscure(*73*). One attribute common to these toxins is that proto-toxin genes are expressed in a variety of tissues. As a result, meta-venom genes likely already interact with ‘future’ toxin genes in other tissues, facilitating their eventual recruitment into the venom.

## Conclusion

When comparing between organisms, it is important to remember that all extant taxa have been evolving for the same amount of time, and that all lineages have experienced different degrees of trait loss and gain(*74*). Therefore, most organisms typically show combinations of both ancestral and derived characters(*75*). Despite being derived phenotypes experiencing strong selection, snake venoms rely on a conserved secretory GRN that is expressed in ancestral saliva and other tissues. Key components of the GRN appear to have been exapted for the evolution of snake and other vertebrate oral venoms. Rather than being non-homologous products of convergent evolution, as previously believed(*38*, *39*, *76*), gene co-expression analysis revealed that these venom systems share a deep homology at the level of regulatory architectures. For example, independent recruitment of kallikreins, most likely represents parallelism rather than constraint. Furthermore, if some animals like mice or rats indeed use salivary proteins as weapons(*58*), which seems plausible given that they use bites during combat for dominance, their saliva would fit common definitions of “venom”(*38*). If so, the evolution of envenomation in vertebrates may be much more common than currently recognized, and the line between vertebrates with and without oral venoms much less clear.

## Additional information

Supplementary code, data, figures, and table can be found at https://github.com/agneeshbarua/Metavenom

## Methods

### RNA extraction and sequencing

RNA was extracted from 30 specimens of *Protobothrops mucrosquamatus* which were collected from various localities throughout Okinawa, Japan. Venom glands were harvested from all 30 specimens while non-venom tissues were harvested from 5 specimens. Specimens had almost equal distribution of male and female (m: 21, f: 26) (Supplementary table M1). Venom was extracted from all specimens at Day 0 and glands were harvested at several time points (Days 1, 2, 4, and 8). RNA-seq libraries were prepared as described in(*77*) and(*10*). Reads were mapped using Bowtie 2 within the RSEM package, which was also used to quantify transcript abundance(*78*). Raw RNA-seq reads are available under NCBI accession PRJDB4386. Further details like specific locations of sampling and generation of RNA data can be found in(*12*).

### Network construction

Weighted gene co-expression analysis was conducted using the *WGCNA* package in R(*23*). The input data consisted of a regularized log transformed matrix of 18,313 genes (as columns) and 29 libraries (as rows) of the venom gland which was filtered for low expressed transcripts (tpm < 0.05). One of the venom gland libraries was excluded in all further analysis due to low spike in (online supplementary, methods section). A characteristic organizational feature of biological networks is a ‘scale-free’ topology, where connections follow a power-law distribution, such that there are very few nodes with very many connections and vice-versa(*79*, *80*). To attain scale-free topology, a soft threshold of 13 was selected based on results from the ‘pickSoftThreshold’ function in the *WGCNA* package (Supplementaryfigure M4). After a soft threshold was estimated, a hierarchical clustering algorithm was used to identify modules of highly connected genes. A threshold of 0.2 and minimum module size = 30 was used to merge very similar expression profiles to obtain a total of 29 modules. We used the ‘modulePreservation’ function to calculate preservation of module characteristics of the meta-venom module, between a reference and test data set. In all cases, the reference data set was the meta-venom module, while the test was a topological overlap matrix (TOM) from either non-venom tissues or venom tissue in cobra. The Zsummay is a composite statistic that combines statistical summaries of network density and connectivity to get a reliable estimate of whether network characteristics are preserved between reference and test(*24*). Simulations revealed that a threshold of 2>Zsummary<10 indicates weak to moderate evidence of preservation, while Zsummary > 10 implies strong preservation, Zsummary < 2 implies no preservation(*24*).

Differential gene expression analysis was carried out in *edgeR(81)*. Transcripts with missing or very low read counts were filtered out before performing the tests. Libraries were normalized (using suggested TMM values) to account for compositional bias as well as account for any size variations between libraries. We performed an analysis of variance (ANOVA)-like test to identify differentially expressed genes between four tissue groups; venom gland, liver, kidney, and heart. A quasi-likelihood F-test was then applied to identify differentially expressed genes between the four groups (at *p* < 0.05 significance). Gene expression plots were made using the same libraries that we used to estimate differential gene expression (at day =1).

#### External validation of module preservation

We conducted an external validation of our data and WGCNA algorithm parameters using an external study of human salivary gland gene expression (*25*). While this data set uses specimens with salivary pathology and was carried out on microarrays. We expected that despite these differences, if the meta-venom is conserved, it will show overlap with one or more modules inferred in the human data. We tested for overlap using Fisher’s exact tests correcting for multiple comparisons using the Benjamini-Hochberg procedure with the false discovery rate set at 0.05.

### Functional annotation of gene sets

GO terms of habu genes were annotated using the free version of Blast2GO software (using a BLAST e-value cutoff ≤ 10^−3^)(*82*). We used both BLAST and InterProt results of the *Protobothrops mucrosquamatus* genome (PRJDB4386) as input for Blast2GO. Using both nucleotide and protein sequences allowed more accurate annotation of GO terms. GO terms enrichment analysis was carried out using the *GOstats* package in R(*83*). Depending on the analysis (eg GO enrichment of meta-venom genes or enrichment of expanded gene families) different gene sets were used as the test data and GO annotations (of the set of all genes) form Blast2GO was used as the ‘universe’.

### Orthology estimate and comparative transcriptomics

Orthologs for habu (*Protobothrops mucrosquamatus* ncbi tax id: 103944), human (*Homo sapiens* 9606), chimp (*Pan troglodytes*: 9598), mouse (*Mus musculus*: 10090), dog (*Canis familiaris*: 9615), anole (*Anolis carolinensis*: 28377), chicken (*Gallus gallus*: 9031), and frog (*Xenopus tropicalis*: 8364) were obtained from the ‘Gene’ database of NCBI (ftp://ftp.ncbi.nlm.nih.gov/gene/DATA/gene_orthologs.gz). These orthologs were calculated by NCBI’s Eukaryotic Genome Annotation pipeline that combines both protein sequence similarity as well as local synteny information. Furthermore, orthologous relations were additionally assigned after manual curation. A combination of command line and R scripts were used to extract a list of one-to-one orthologs shared between all 8 taxa (Online supplementary material). In addition to using the orthologs defined by NCBI, we carried out phylogenetic ortholog estimation using OrthoFinder (OF)(*28*). OF uses protein sequences to form orthogroups and then combines information from gene trees and species trees to distinguish between gene copies arising from speciation or duplication events within lineages. OF also has the added advantage of removing any errors that tend to occur during similarity based assignment of orthologs(*84*). Protein sequences for 8 taxa were obtained from Ensembl(*85*). Cobra (*Naja naja*: 35670) protein and transcript sequences were obtained by request from the authors of(*29*). Using both these approaches we obtained a combined list of 2682 one-to-one expressed orthologs (see next section) between 9 taxa. From these we filtered meta-venom orthologs based on the habu genes present in the meta-venom. This results in a list of 546 expressed meta-venom orthologs found in all 9 taxa.

RNA data for each species and tissue were obtained from the SRA database (Supplementary table M2). Datasets were from a variety of sources including published studies(*29*, *86*–*89*) and large scale sequencing projects like the Broad Institute Canine Genomic Resources and the ENCODE project (*90*). Where possible, at least 3 libraries for each tissue from each taxa were used to compile our comparative dataset, and only data generated from healthy, adult tissues was used. We used the ‘fasterq-dump’ function in *SRA toolkit 2.9.1* (https://github.com/ncbi/sra-tools/wiki) to download fastq files, which were quantified using

*kallisto* (*91*). *Kallisto* indices for human, mouse, chimp, dog, anole, frog, and chicken were created using GTF and cDNA files from the Ensembl database(*85*). Index for cobra was made using annotation and transcript files from(*29*). For single end reads we set length parameter to 350 and standard deviation of length fragment to 150. A custom R script was used to aggregate transcript-level read counts to gene-level read counts.. Once total tpm was obtained for each tissue from each taxa, the data were filtered to obtain a final dataset of one-to-one orthologs expressed across all tissues across all 9 taxa. To allow for comparisons across samples, expression levels were normalized. Normalization was carried out by adding a pseudo count of 1 × 10^−5^ (to prevent log[0] scores), followed by log2 transformation. The transformed data was then quantile normalized among samples. Quantile normalization ensured equal across sample distribution of gene expression levels so as to minimize the effects of technical artefacts(*92*, *93*).

Our aim was to identify any conserved pattern of expression present between homologous tissues from multiple taxa, however identifying patterns in expression data from multiple species as well as multiple studies requires the removal of their respective batch effects(*94*). The batch effect imparted by species is due to the level of shared functionality of genetic processes, where evolutionary changes (during speciation) in shared molecular machinery will simultaneously alter the expression of genes in all tissues, thereby masking any historical signals of homology(*95*, *96*). To remove these batch effects and identify patterns (if any) of homology in expression between tissues we used an empirical Bayes method (implemented via the ComBat function in the *sva* R package)(*97*). We used the plotPCA function in the *DESeq2* package(*98*) to carry out principal component analysis. Using both species and study as batch effects produced similar results, although species explained more variation and provided better resolution of underlying tissue specific trends (online supplementary material).

### Gene family evolution

Gene family evolution across amniotes was investigated using CAFE v5.0(*34*, *99*). CAFE models gene family evolution across a species tree using a stochastic birth and death process. An ultrametric species tree was drawn in Mesquite(*100*) and divergence times were estimated using http://www.timetree.org/. Protein sequences for 7 taxa were obtained from Ensembl the rest (habu and cobra) from NCBI. Gene families were inferred with BLAST and MCL (implemented in CAFE), using proteins present in the meta-venom as query sequences. This resulted in 250 estimated gene families. Although most of our taxa are model organisms with well assembled genomes, for increased statistical robustness, we estimated an error model due to genome assembly error which was later used for λ analysis(*33*)(Supplementary table M3). The rate parameter λ describes the probability that any gene will either be gained or lost, where a higher λ denotes rapid gene family evolution (*34*). We used a global λ (λ1) as our null model and a different rate parameter (λ2) for the lineage leading upto venomous snakes to test the hypothesis that gene families evolved faster in the lineage leading up to venomous snakes compared to other lineages. Simulations of gene families from observed data and a subsequent likelihood ratio test using the global λ (λ1) estimate and lineage specific λ (λ2) was used to determine significance. Once the log-likelihoods were obtained, lhtest.R script (provided by CAFE) was used to create histogram with a null distribution obtained from simulations. Significance is determined by how far left the observed likelihood ratio (2 x lnLglobal - lnLmulti) would fall on the tail of the distribution. In our case the likelihood ratio count would fall on the far left of the distribution indicating a very low *p* value (Supplementary figure M5). Along with inferring rates of gene family evolution, CAFE also determines expansions or contractions in gene size by calculating ancestral states at nodes along the tree. For each gene family CAFE computes a *p*-value associated with the gene family size in extant species given the model of gene family evolution(*99*). This was used to determine which gene families underwent significant expansion, contraction, or stayed the same in venomous snakes (Supplementary table 5).

## References

1. T. F. Duda Jr, S. R. Palumbi, Evolutionary diversification of multigene families: allelic selection of toxins in predatory cone snails. Mol. Biol. Evol. 17, 1286–1293 (2000).

2. Y. Moran, H. Weinberger, J. C. Sullivan, A. M. Reitzel, J. R. Finnerty, M. Gurevitz, Concerted evolution of sea anemone neurotoxin genes is revealed through analysis of the Nematostella vectensis genome. Mol. Biol. Evol. 25, 737–747 (2008).

3. D. R. Rokyta, A. R. Lemmon, M. J. Margres, K. Aronow, The venom-gland transcriptome of the eastern diamondback rattlesnake (Crotalus adamanteus). BMC Genomics. 13, 312 (2012).

4. D. R. Rokyta, K. P. Wray, M. J. Margres, The genesis of an exceptionally lethal venom in the timber rattlesnake (Crotalus horridus) revealed through comparative venom-gland transcriptomics. BMC Genomics. 14, 394 (2013).

5. J. M. Surm, H. L. Smith, B. Madio, E. A. B. Undheim, G. F. King, B. R. Hamilton, C. A. van der Burg, A. Pavasovic, P. J. Prentis, A process of convergent amplification and tissue-specific expression dominates the evolution of toxin and toxin-like genes in sea anemones. Mol. Ecol. 28, 2272–2289 (2019).

6. M. J. Margres, K. P. Wray, A. T. B. Hassinger, M. J. Ward, J. J. McGivern, E. M. Lemmon, A. R. Lemmon, D. R. Rokyta, Quantity, not quality: rapid adaptation in a polygenic trait proceeded exclusively through expression differentiation. Mol. Biol. Evol. 34, 3099–3110 (2017).

7. A. Barua, A. S. Mikheyev, Many Options, Few Solutions: Over 60 Million Snakes Converged on a Few Optimal Venom Formulations. Mol. Biol. Evol. 36, 1964–1974 (2019).

8. V. Schendel, L. D. Rash, R. A. Jenner, E. A. B. Undheim, The Diversity of Venom: The Importance of Behavior and Venom System Morphology in Understanding Its Ecology and Evolution. Toxins. 11, 666 (2019).

9. G. Zancolli, N. R. Casewell, Venom systems as models for studying the origin and regulation of evolutionary novelties. Mol. Biol. Evol. (2020), doi:10.1093/molbev/msaa133.

10. S. D. Aird, S. Aggarwal, A. Villar-Briones, M. M.-Y. Tin, K. Terada, A. S. Mikheyev, Snake venoms are integrated systems, but abundant venom proteins evolve more rapidly. BMC Genomics. 16, 647 (2015).

11. A. Barua, A. S. Mikheyev, Toxin expression in snake venom evolves rapidly with constant shifts in evolutionary rates. Proceedings of the Royal Society B: Biological Sciences. 287, 20200613 (2020).

12. S. D. Aird, J. Arora, A. Barua, L. Qiu, K. Terada, A. S. Mikheyev, Population genomic analysis of a pitviper reveals microevolutionary forces underlying venom chemistry. Genome Biol. Evol. 9, 2640–2649 (2017).

13. A. Malhotra, S. Creer, J. B. Harris, R. S. Thorpe, The importance of being genomic: Non-coding and coding sequences suggest different models of toxin multi-gene family evolution. Toxicon. 107, 344–358 (2015).

14. B. G. Fry, From genome to “venome”: molecular origin and evolution of the snake venom proteome inferred from phylogenetic analysis of toxin sequences and related body …. Genome Res. (2005) (available at http://genome.cshlp.org/content/15/3/403.short).

15. B. G. Fry, H. Scheib, L. van der Weerd, B. Young, J. McNaughtan, S. F. R. Ramjan, N. Vidal, R. E. Poelmann, J. A. Norman, Evolution of an arsenal: structural and functional diversification of the venom system in the advanced snakes (Caenophidia). Mol. Cell. Proteomics. 7, 215–246 (2008).

16. A.-L. Barabási, Z. N. Oltvai, Network biology: understanding the cell’s functional organization. Nat. Rev. Genet. 5, 101–113 (2004).

17. J. A. Miller, S. Horvath, D. H. Geschwind, Divergence of human and mouse brain transcriptome highlights Alzheimer disease pathways. Proc. Natl. Acad. Sci. U. S. A. 107, 12698–12703 (2010).

18. M. C. Oldham, S. Horvath, D. H. Geschwind, Conservation and evolution of gene coexpression networks in human and chimpanzee brains. Proc. Natl. Acad. Sci. U. S. A. 103, 17973–17978 (2006).

19. M. Filteau, S. A. Pavey, J. St-Cyr, L. Bernatchez, Gene coexpression networks reveal key drivers of phenotypic divergence in lake whitefish. Mol. Biol. Evol. 30, 1384–1396 (2013).

20. J. M. Stuart, E. Segal, D. Koller, S. K. Kim, A gene-coexpression network for global discovery of conserved genetic modules. Science. 302, 249–255 (2003).

21. J. D. Allen, Y. Xie, M. Chen, L. Girard, G. Xiao, Comparing statistical methods for constructing large scale gene networks. PLoS One. 7, e29348 (2012).

22. V. A. H.-T. Guido Sanguinetti, Ed., Gene Regulatory Networks.pdf (Springer Science and Business Media, 233 Spring Street, New York, NY 10013, USA, 2019), Springer Protocols.

23. P. Langfelder, S. Horvath, WGCNA: an R package for weighted correlation network analysis. BMC Bioinformatics. 9, 559 (2008).

24. P. Langfelder, R. Luo, M. C. Oldham, S. Horvath, Is my network module preserved and reproducible? PLoS Comput. Biol. 7, e1001057 (2011).

25. S. Horvath, A. N. M. Nazmul-Hossain, R. P. E. Pollard, F. G. M. Kroese, A. Vissink, C. G. M. Kallenberg, F. K. L. Spijkervet, H. Bootsma, S. A. Michie, S. U. Gorr, A. B. Peck, C. Cai, H. Zhou, D. T. W. Wong, Systems analysis of primary Sjögren’s syndrome pathogenesis in salivary glands identifies shared pathways in human and a mouse model. Arthritis Res. Ther. 14, R238 (2012).

26. D. Ron, P. Walter, Signal integration in the endoplasmic reticulum unfolded protein response. Nat. Rev. Mol. Cell Biol. 8, 519–529 (2007).

27. NCBI Resource Coordinators, Database resources of the National Center for Biotechnology Information. Nucleic Acids Res. 44, D7–19 (2016).

28. D. M. Emms, S. Kelly, OrthoFinder: phylogenetic orthology inference for comparative genomics. Genome Biol. 20, 238 (2019).

29. K. Suryamohan, S. P. Krishnankutty, J. Guillory, M. Jevit, M. S. Schröder, M. Wu, B. Kuriakose, O. K. Mathew, R. C. Perumal, I. Koludarov, L. D. Goldstein, K. Senger, M. D. Dixon, D. Velayutham, D. Vargas, S. Chaudhuri, M. Muraleedharan, R. Goel, Y.-J. J. Chen, A. Ratan, P. Liu, B. Faherty, G. de la Rosa, H. Shibata, M. Baca, M. Sagolla, J. Ziai, G. A. Wright, D. Vucic, S. Mohan, A. Antony, J. Stinson, D. S. Kirkpatrick, R. N. Hannoush, S. Durinck, Z. Modrusan, E. W. Stawiski, K. Wiley, T. Raudsepp, R. M. Kini, A. Zachariah, S. Seshagiri, The Indian cobra reference genome and transcriptome enables comprehensive identification of venom toxins. Nat. Genet. 52, 106–117 (2020).

30. A. Breschi, S. Djebali, J. Gillis, D. D. Pervouchine, A. Dobin, C. A. Davis, T. R. Gingeras, R. Guigó, Gene-specific patterns of expression variation across organs and species. Genome Biol. 17, 151 (2016).

31. N. L. Dowell, M. W. Giorgianni, V. A. Kassner, J. E. Selegue, E. E. Sanchez, S. B. Carroll, The deep origin and recent loss of venom toxin genes in rattlesnakes. Curr. Biol. 26, 2434–2445 (2016).

32. M. W. Giorgianni, N. L. Dowell, S. Griffin, V. A. Kassner, J. E. Selegue, S. B. Carroll, The origin and diversification of a novel protein family in venomous snakes. Proc. Natl. Acad. Sci. U. S. A. 117, 10911–10920 (2020).

33. M. V. Han, G. W. C. Thomas, J. Lugo-Martinez, M. W. Hahn, Estimating gene gain and loss rates in the presence of error in genome assembly and annotation using CAFE 3. Mol. Biol. Evol. 30, 1987–1997 (2013).

34. M. W. Hahn, T. De Bie, J. E. Stajich, C. Nguyen, N. Cristianini, Estimating the tempo and mode of gene family evolution from comparative genomic data. Genome Res. 15, 1153–1160 (2005).

35. M. R. J. Carlson, B. Zhang, Z. Fang, P. S. Mischel, S. Horvath, S. F. Nelson, Gene connectivity, function, and sequence conservation: predictions from modular yeast co-expression networks. BMC Genomics. 7, 40 (2006).

36. M. Rotival, E. Petretto, Leveraging gene co-expression networks to pinpoint the regulation of complex traits and disease, with a focus on cardiovascular traits. Brief. Funct. Genomics. 13, 66–78 (2014).

37. A. J. Mason, M. J. Margres, J. L. Strickland, D. R. Rokyta, M. Sasa, C. L. Parkinson, Trait differentiation and modular toxin expression in palm-pitvipers. BMC Genomics. 21, 147 (2020).

38. R. Ligabue-Braun, H. Verli, C. R. Carlini, Venomous mammals: a review. Toxicon. 59, 680–695 (2012).

39. N. R. Casewell, D. Petras, D. C. Card, V. Suranse, A. M. Mychajliw, D. Richards, I. Koludarov, L.-O. Albulescu, J. Slagboom, B.-F. Hempel, N. M. Ngum, R. J. Kennerley, J. L. Brocca, G. Whiteley, R. A. Harrison, F. M. S. Bolton, J. Debono, F. J. Vonk, J. Alföldi, J. Johnson, E. K. Karlsson, K. Lindblad-Toh, I. R. Mellor, R. D. Süssmuth, B. G. Fry, S. Kuruppu, W. C. Hodgson, J. Kool, T. A. Castoe, I. Barnes, K. Sunagar, E. A. B. Undheim, S. T. Turvey, Solenodon genome reveals convergent evolution of venom in eulipotyphlan mammals. Proc. Natl. Acad. Sci. U. S. A. 116, 25745–25755 (2019).

40. E. B. Rosenblum, C. E. Parent, E. E. Brandt, The molecular basis of phenotypic convergence. Annu. Rev. Ecol. Evol. Syst. 45, 203–226 (2014).

41. S. P. Mackessy, A. J. Saviola, Understanding Biological Roles of Venoms Among the Caenophidia: The Importance of Rear-Fanged Snakes. Integr. Comp. Biol. 56, 1004–1021 (2016).

42. G. Xu, S. R. Jaffrey, The new landscape of protein ubiquitination. Nat. Biotechnol. 29, 1098–1100 (2011).

43. J. Hwang, L. Qi, Quality Control in the Endoplasmic Reticulum: Crosstalk between ERAD and UPR pathways. Trends Biochem. Sci. 43, 593–605 (2018).

44. M. Michalak, J. Groenendyk, E. Szabo, L. I. Gold, M. Opas, Calreticulin, a multi-process calcium-buffering chaperone of the endoplasmic reticulum. Biochem. J. 417, 651–666 (2009).

45. T. J. Bergmann, I. Fregno, F. Fumagalli, A. Rinaldi, F. Bertoni, P. J. Boersema, P. Picotti, M. Molinari, Chemical stresses fail to mimic the unfolded protein response resulting from luminal load with unfolded polypeptides. J. Biol. Chem. 293, 5600–5612 (2018).

46. M. Bai, R. Vozdek, A. Hnízda, C. Jiang, B. Wang, L. Kuchar, T. Li, Y. Zhang, C. Wood, L. Feng, Y. Dang, D. K. Ma, Conserved roles of C. elegans and human MANFs in sulfatide binding and cytoprotection. Nat. Commun. 9, 897 (2018).

47. H. Safavi-Hemami, Q. Li, R. L. Jackson, A. S. Song, W. Boomsma, P. K. Bandyopadhyay, C. W. Gruber, A. W. Purcell, M. Yandell, B. M. Olivera, L. Ellgaard, Rapid expansion of the protein disulfide isomerase gene family facilitates the folding of venom peptides. Proc. Natl. Acad. Sci. U. S. A. 113, 3227–3232 (2016).

48. L. Ellgaard, L. W. Ruddock, The human protein disulphide isomerase family: substrate interactions and functional properties. EMBO Rep. 6, 28–32 (2005).

49. M. Michalak, E. F. Corbett, N. Mesaeli, K. Nakamura, M. Opas, Calreticulin: one protein, one gene, many functions. Biochem. J. 344 Pt 2, 281–292 (1999).

50. D. T. Rutkowski, R. S. Hegde, Regulation of basal cellular physiology by the homeostatic unfolded protein response. J. Cell Biol. 189, 783–794 (2010).

51. X. Yang, K. Matsuda, P. Bialek, S. Jacquot, H. C. Masuoka, T. Schinke, L. Li, S. Brancorsini, P. Sassone-Corsi, T. M. Townes, A. Hanauer, G. Karsenty, ATF4 Is a Substrate of RSK2 and an Essential Regulator of Osteoblast Biology. Cell. 117(2004), pp. 387–398.

52. A.-H. Lee, E. F. Scapa, D. E. Cohen, L. H. Glimcher, Regulation of hepatic lipogenesis by the transcription factor XBP1. Science. 320, 1492–1496 (2008).

53. Y. Wang, L. Vera, W. H. Fischer, M. Montminy, The CREB coactivator CRTC2 links hepatic ER stress and fasting gluconeogenesis. Nature. 460, 534–537 (2009).

54. D. Senft, Z. A. Ronai, UPR, autophagy, and mitochondria crosstalk underlies the ER stress response. Trends Biochem. Sci. 40, 141–148 (2015).

55. A. Pavlopoulou, G. Pampalakis, I. Michalopoulos, G. Sotiropoulou, Evolutionary history of tissue kallikreins. PLoS One. 5, e13781 (2010).

56. V. L. Koumandou, A. Scorilas, Evolution of the plasma and tissue kallikreins, and their alternative splicing isoforms. PLoS One. 8, e68074 (2013).

57. R. C. Karn, C. M. Laukaitis, Positive selection shaped the convergent evolution of independently expanded kallikrein subfamilies expressed in mouse and rat saliva proteomes. PLoS One. 6, e20979 (2011).

58. M. Hiramatsu, K. Hatakeyama, N. Minami, Male mouse submaxillary gland secretes highly toxic proteins. Experientia. 36, 940–942 (1980).

59. D. H. Dean, R. N. Hiramoto, Lethal effect of male rat submandibular gland homogenate for rat neonates. J. Oral Pathol. 14, 666–669 (1985).

60. J. C. Huang, K. Hoshino, Y. T. Kim, F. S. Chebib, Species and strain differences in the lethal factor of the mouse submandibular gland. Can. J. Physiol. Pharmacol. 55, 1107–1111 (1977).

61. M. Kita, Y. Nakamura, Y. Okumura, S. D. Ohdachi, Y. Oba, M. Yoshikuni, H. Kido, D. Uemura, Blarina toxin, a mammalian lethal venom from the short-tailed shrew Blarina brevicauda: Isolation and characterization. Proc. Natl. Acad. Sci. U. S. A. 101, 7542–7547 (2004).

62. G. Datta, A. T. Tu, Structure and other chemical characterizations of gila toxin, a lethal toxin from lizard venom. J. Pept. Res. 50, 443–450 (2009).

63. P. L. Mailho-Fontana, M. M. Antoniazzi, C. Alexandre, D. C. Pimenta, J. M. Sciani, E. D. Brodie Jr, C. Jared, Morphological Evidence for an Oral Venom System in Caecilian Amphibians. iScience, 101234 (2020).

64. S. Vaiyapuri, K. Sunagar, J. M. Gibbins, T. N. W. Jackson, T. Reeks, B. G. Fry, Kallikrein enzymes. Venomous Reptiles and Their Toxins: Evolution, Pathophysiology and Biodiscovery; Fry, BG, Ed, 267–280 (2015).

65. S. D. Aird, Ophidian envenomation strategies and the role of purines. Toxicon. 40, 335–393 (2002).

66. K. Sunagar, W. E. Johnson, S. J. O’Brien, V. Vasconcelos, A. Antunes, Evolution of CRISPs associated with toxicoferan-reptilian venom and mammalian reproduction. Mol. Biol. Evol. 29, 1807–1822 (2012).

67. W. H. Heyborne, S. P. Mackessy, in Handbook of Venoms and Toxins of Reptiles, S. P. Mackessy, Ed. (CRC Press, 2016), pp. 325–335.

68. R. Kakumanu, W. C. Hodgson, R. Ravi, A. Alagon, R. J. Harris, A. Brust, P. F. Alewood, B. K. Kemp-Harper, B. G. Fry, Vampire Venom: Vasodilatory Mechanisms of Vampire Bat (Desmodus rotundus) Blood Feeding. Toxins. 11 (2019), doi:10.3390/toxins11010026.

69. J. Ekström, R. Ekman, R. Håkanson, S. Sjögren, F. Sundler, Calcitonin gene-related peptide in rat salivary glands: neuronal localization, depletion upon nerve stimulation, and effects on salivation in relation to substance P. Neuroscience. 26, 933–949 (1988).

70. S. D. Brain, T. J. Williams, J. R. Tippins, H. R. Morris, I. MacIntyre, Calcitonin gene-related peptide is a potent vasodilator. Nature. 313, 54–56 (1985).

71. F. J. Vonk, N. R. Casewell, C. V. Henkel, A. M. Heimberg, H. J. Jansen, R. J. R. McCleary, H. M. E. Kerkkamp, R. A. Vos, I. Guerreiro, J. J. Calvete, W. Wüster, A. E. Woods, J. M. Logan, R. A. Harrison, T. A. Castoe, A. P. J. de Koning, D. D. Pollock, M. Yandell, D. Calderon, C. Renjifo, R. B. Currier, D. Salgado, D. Pla, L. Sanz, A. S. Hyder, J. M. C. Ribeiro, J. W. Arntzen, G. E. E. J. M. van den Thillart, M. Boetzer, W. Pirovano, R. P. Dirks, H. P. Spaink, D. Duboule, E. McGlinn, R. M. Kini, M. K. Richardson, The king cobra genome reveals dynamic gene evolution and adaptation in the snake venom system. Proc. Natl. Acad. Sci. U. S. A. 110, 20651–20656 (2013).

72. J. A. Jury, A. C. Perry, L. Hall, Identification, sequence analysis and expression of transcripts encoding a putative metalloproteinase, eMDC II, in human and macaque epididymis. Mol. Hum. Reprod. 5, 1127–1134 (1999).

73. I. Koludarov, T. N. W. Jackson, A. Pozzi, A. S. Mikheyev, Family saga: reconstructing the evolutionary history of a functionally diverse gene family reveals complexity at the genetic origins of novelty. bioRxiv (2019) (available at https://www.biorxiv.org/content/10.1101/583344v1.abstract).

74. R. R. Strathmann, D. J. Eernisse, What Molecular Phylogenies Tell Us about the Evolution of Larval Forms. Integr. Comp. Biol. 34, 502–512 (1994).

75. R. A. Jenner, Unburdening evo-devo: ancestral attractions, model organisms, and basal baloney. Dev. Genes Evol. 216, 385–394 (2006).

76. K. E. Folinsbee, Evolution of venom across extant and extinct eulipotyphlans. C. R. Palevol. 12, 531–542 (2013).

77. S. D. Aird, Y. Watanabe, A. Villar-Briones, M. C. Roy, K. Terada, A. S. Mikheyev, Quantitative high-throughput profiling of snake venom gland transcriptomes and proteomes (Ovophis okinavensis and Protobothrops flavoviridis). BMC Genomics. 14, 790 (2013).

78. B. Langmead, S. L. Salzberg, Fast gapped-read alignment with Bowtie 2. Nat. Methods. 9, 357–359 (2012).

79. B. Zhang, S. Horvath, A general framework for weighted gene co-expression network analysis. Stat. Appl. Genet. Mol. Biol. 4, Article17 (2005).

80. M. L. Siegal, D. E. L. Promislow, A. Bergman, Functional and evolutionary inference in gene networks: does topology matter? Genetica. 129, 83–103 (2007).

81. M. D. Robinson, D. J. McCarthy, G. K. Smyth, edgeR: a Bioconductor package for differential expression analysis of digital gene expression data. Bioinformatics. 26, 139–140 (2010).

82. S. Götz, J. M. García-Gómez, J. Terol, T. D. Williams, S. H. Nagaraj, M. J. Nueda, M. Robles, M. Talón, J. Dopazo, A. Conesa, High-throughput functional annotation and data mining with the Blast2GO suite. Nucleic Acids Res. 36, 3420–3435 (2008).

83. S. Falcon, R. Gentleman, Using GOstats to test gene lists for GO term association. Bioinformatics. 23, 257–258 (2007).

84. D. M. Emms, S. Kelly, OrthoFinder: solving fundamental biases in whole genome comparisons dramatically improves orthogroup inference accuracy. Genome Biol. 16, 157 (2015).

85. A. D. Yates, P. Achuthan, W. Akanni, J. Allen, J. Allen, J. Alvarez-Jarreta, M. R. Amode, I. M. Armean, A. G. Azov, R. Bennett, J. Bhai, K. Billis, S. Boddu, J. C. Marugán, C. Cummins, C. Davidson, K. Dodiya, R. Fatima, A. Gall, C. G. Giron, L. Gil, T. Grego, L. Haggerty, E. Haskell, T. Hourlier, O. G. Izuogu, S. H. Janacek, T. Juettemann, M. Kay, I. Lavidas, T. Le, D. Lemos, J. G. Martinez, T. Maurel, M. McDowall, A. McMahon, S. Mohanan, B. Moore, M. Nuhn, D. N. Oheh, A. Parker, A. Parton, M. Patricio, M. P. Sakthivel, A. I. Abdul Salam, B. M. Schmitt, H. Schuilenburg, D. Sheppard, M. Sycheva, M. Szuba, K. Taylor, A. Thormann, G. Threadgold, A. Vullo, B. Walts, A. Winterbottom, A. Zadissa, M. Chakiachvili, B. Flint, A. Frankish, S. E. Hunt, G. IIsley, M. Kostadima, N. Langridge, J. E. Loveland, F. J. Martin, J. Morales, J. M. Mudge, M. Muffato, E. Perry, M. Ruffier, S. J. Trevanion, F. Cunningham, K. L. Howe, D. R. Zerbino, P. Flicek, Ensembl 2020. Nucleic Acids Res. 48, D682–D688 (2020).

86. D. Brawand, M. Soumillon, A. Necsulea, P. Julien, G. Csárdi, P. Harrigan, M. Weier, A. Liechti, A. Aximu-Petri, M. Kircher, F. W. Albert, U. Zeller, P. Khaitovich, F. Grützner, S. Bergmann, R. Nielsen, S. Pääbo, H. Kaessmann, The evolution of gene expression levels in mammalian organs. Nature. 478, 343–348 (2011).

87. N. L. Barbosa-Morais, M. Irimia, Q. Pan, H. Y. Xiong, S. Gueroussov, L. J. Lee, V. Slobodeniuc, C. Kutter, S. Watt, R. Colak, T. Kim, C. M. Misquitta-Ali, M. D. Wilson, P. M. Kim, D. T. Odom, B. J. Frey, B. J. Blencowe, The evolutionary landscape of alternative splicing in vertebrate species. Science. 338, 1587–1593 (2012).

88. R. Marin, D. Cortez, F. Lamanna, M. M. Pradeepa, E. Leushkin, P. Julien, A. Liechti, J. Halbert, T. Brüning, K. Mössinger, T. Trefzer, C. Conrad, H. N. Kerver, J. Wade, P. Tschopp, H. Kaessmann, Convergent origination of a Drosophila-like dosage compensation mechanism in a reptile lineage. Genome Res. 27, 1974–1987 (2017).

89. M. Alame, E. Cornillot, V. Cacheux, G. Tosato, M. Four, L. De Oliveira, S. Gofflot, P. Delvenne, E. Turtoi, S. Cabello-Aguilar, M. Nishiyama, A. Turtoi, V. Costes-Martineau, J. Colinge, The molecular landscape and microenvironment of salivary duct carcinoma reveal new therapeutic opportunities. Theranostics. 10, 4383–4394 (2020).

90. ENCODE Project Consortium, A user’s guide to the encyclopedia of DNA elements (ENCODE). PLoS Biol. 9, e1001046 (2011).

91. N. L. Bray, H. Pimentel, P. Melsted, L. Pachter, Near-optimal probabilistic RNA-seq quantification. Nat. Biotechnol. 34, 525–527 (2016).

92. B. M. Bolstad, R. A. Irizarry, M. Astrand, T. P. Speed, A comparison of normalization methods for high density oligonucleotide array data based on variance and bias. Bioinformatics. 19, 185–193 (2003).

93. B. Qiu, R. S. Larsen, N.-C. Chang, J. Wang, J. J. Boomsma, G. Zhang, Towards reconstructing the ancestral brain gene-network regulating caste differentiation in ants. Nat Ecol Evol. 2, 1782–1791 (2018).

94. Y. Gilad, O. Mizrahi-Man, A reanalysis of mouse ENCODE comparative gene expression data. F1000Res. 4, 121 (2015).

95. J. M. Musser, G. P. Wagner, Character trees from transcriptome data: Origin and individuation of morphological characters and the so-called “species signal”: TRANSCRIPTOME-BASED CHARACTER TREES. J. Exp. Zool. 324, 588–604 (2015).

96. C. Liang, J. M. Musser, A. Cloutier, R. O. Prum, G. P. Wagner, Pervasive Correlated Evolution in Gene Expression Shapes Cell and Tissue Type Transcriptomes. Genome Biol. Evol. 10, 538–552 (2018).

97. J. T. Leek, W. E. Johnson, H. S. Parker, A. E. Jaffe, J. D. Storey, The sva package for removing batch effects and other unwanted variation in high-throughput experiments. Bioinformatics. 28, 882–883 (2012).

98. M. I. Love, W. Huber, S. Anders, Moderated estimation of fold change and dispersion for RNA-seq data with DESeq2. Genome Biol. 15, 550 (2014).

99. T. De Bie, N. Cristianini, J. P. Demuth, M. W. Hahn, CAFE: a computational tool for the study of gene family evolution. Bioinformatics. 22, 1269–1271 (2006).

100. W. P. A. D. R. M. Maddison, Mesquite: a modular system for evolutionary analysis. Version 3.61 (2019), (available at http://www.mesquiteproject.org).

